# A low-power communication scheme for wireless, 1000 channel brain-machine interfaces

**DOI:** 10.1101/2022.03.11.483996

**Authors:** Joseph T. Costello, Samuel R. Nason, Hyochan An, Jungho Lee, Matthew J. Mender, Hisham Temmar, Dylan M. Wallace, Jongyup Lim, Matthew S. Willsey, Parag G. Patil, Taekwang Jang, Jamie D. Phillips, Hun-Seok Kim, David Blaauw, Cynthia A. Chestek

**Affiliations:** Department of Electrical and Computer Engineering, Ann Arbor, MI, USA; Department of Biomedical Engineering, Ann Arbor, MI, USA; Robotics Institute, University of Michigan, Ann Arbor, MI, USA; Department of Neurosurgery, University of Michigan Medical School, Ann Arbor, MI; Department of Information Technology and Electrical Engineering, ETH Zurich, Zurich, Switzerland; Department of Electrical and Computer Engineering, University of Delaware, Newark, DE, USA

**Keywords:** brain-machine interface, neural interface, low-power, wireless communication

## Abstract

**Objective:** Brain-machine interfaces (BMIs) have the potential to restore motor function but are currently limited by electrode count and long-term recording stability. These challenges may be solved through the use of free-floating “motes” which wirelessly transmit recorded neural signals, if power consumption can be kept within safe levels when scaling to thousands of motes. Here, we evaluated a pulse-interval modulation (PIM) communication scheme for infrared (IR)-based motes that aims to reduce the wireless data rate and system power consumption.

**Approach:** To test PIM’s ability to efficiently communicate neural information, we simulated the communication scheme in a real-time closed-loop BMI with non-human primates. Additionally, we performed circuit simulations of an IR-based 1000-mote system to calculate communication accuracy and total power consumption.

**Main Results:** We found that PIM at 1kb/s per channel maintained strong correlations with true firing rate and matched online BMI performance of a traditional wired system. Closed-loop BMI tests suggest that lags as small as 30 ms can have significant performance effects. Finally, unlike other IR communication schemes, PIM is feasible in terms of power, and neural data can accurately be recovered on a receiver using 3mW for 1000 channels.

**Significance:** These results suggest that PIM-based communication could significantly reduce power usage of wireless motes to enable higher channel-counts for high-performance BMIs.

## Introduction

Brain machine interfaces (BMIs) have shown potential for restoring movement and communication to those who suffer from spinal cord injury. BMIs estimate user intentions by recording from electrodes implanted in cortical regions and processing neural data with a decoding algorithm. These systems have allowed participants to control prosthetic arms [1], [2], write text [3], and functionally stimulate paralyzed limbs [4]. Current BMIs in humans use wired electrode arrays, most commonly the Utah array [5], [6]. While Utah arrays have enabled some success in decoding intentions, their performance is still significantly below able-bodied control; part of this performance drop may be attributed to the limited number of electrodes (up to 256 channels, [7]) and low neuronal yield [8], [9]. While multiple Utah arrays can be implanted, it is infeasible to implant tens of arrays in a single cortical region, which would be required for hundreds to thousands of tuned channels. Thus, recent efforts have looked toward other electrode technologies for increasing channel counts and improving chronic recording stability to improve performance of intracortical BMIs.

To increase the number of recording channels, several groups have developed rigid electrode arrays with thousands of channels: The Argo [10] can record from 65k channels using a microwire bundle with 60-300 μm pitch. Neuropixels 2.0 [11] has up to 10k channels on an implant at a 15 μm pitch. While high channel counts could enable better BMI control, these systems have several limitations, which make them difficult to use for chronic motor BMI. First, with thousands of wires, the interconnect for these systems requires large, chronically open holes in the dura mater which may become failure points. This rigid connectorization may also experience micromotion relative to the brain, which may cause unstable recordings, glial scarring and cell loss near the electrode. From the perspective of BMI performance, these high-density electrodes have a large number of recording sites spaced closely together. Therefore, they record from small brain areas, which may have highly correlated neural activity [12]; dispersing channels across a wider region could record from cells with more independent information, allowing for better decoding across more degrees of freedom [13].

Significant progress has been made in developing thin, flexible electrodes that aim to move with the brain and reduce tissue damage. These include the mesh probe [14], which achieves better biocompatibility through a flexible 3D mesh, the Neural Matrix [15], a chronic surface array with over 1000 channels, and a polymer-based microelectrode array with 512 channels [16]. Notably, Neuralink has developed a multi-thousand channel recording system using polymer “thread” electrodes connected a custom recording ASIC [17]. As the channel count of flexible systems increases, however, the wire interconnects must become increasingly unwieldy, resulting in additional failure points. Despite potentially minimal tissue damage, soft electrodes still require large, chronic openings in the dura mater for the wire interconnect.

Alternatively, wireless dust [18], [19] or “motes” show promise in solving the challenges of recording stability and high channel counts, without being limited by the interconnect. Each mote receives wireless power, records single-channel neural data, and transmits data to a receiver. Since these devices are free-floating, they can move with the brain for reduced tissue scarring and can be covered by dura, thereby reducing the risk of cerebrospinal fluid leak and infection. They can be implanted in any desired configuration across large regions. However, a primary challenge with wireless motes is minimizing power usage to limit tissue heating to safe levels, while supporting communication with up to thousands of motes. Several wireless power transfer and communication modalities have been previously explored: The original Neural Dust [18], [19] is a sub-mm mote powered through ultrasound that can be implanted as deep as 5 cm [20]. The Neurograin [21] is powered through a radiofrequency (RF) link and could theoretically support up to 770 devices. Finally, the OWIC [22] and Michigan Mote (Figure 1a, [23], [24]) are powered through infrared (IR) light. Other power modalities and designs have been proposed for stimulation motes, but their power requirements are significantly higher. With each power modality there is a tradeoff between maximum device power and device size, and unique considerations with respect to energy harvesting efficiency and communications methods with device size scaling. Motes using IR power transfer and communications are particularly promising because they can be nearly an order of magnitude smaller than RF and ultrasound-based motes, but have the lowest available power due to the reduced mote area [25](Robinson 2021). For example, benchtop and acute testing has demonstrated that IR motes can be smaller than 250 μm maximum dimension [22], [24] but are then limited to 1.5 μW per device [26]. Thus, IR power may be advantageous for recording-based BMIs that require high channel density spread out across a larger area, if power constraints can be met.

**Figure 1:**
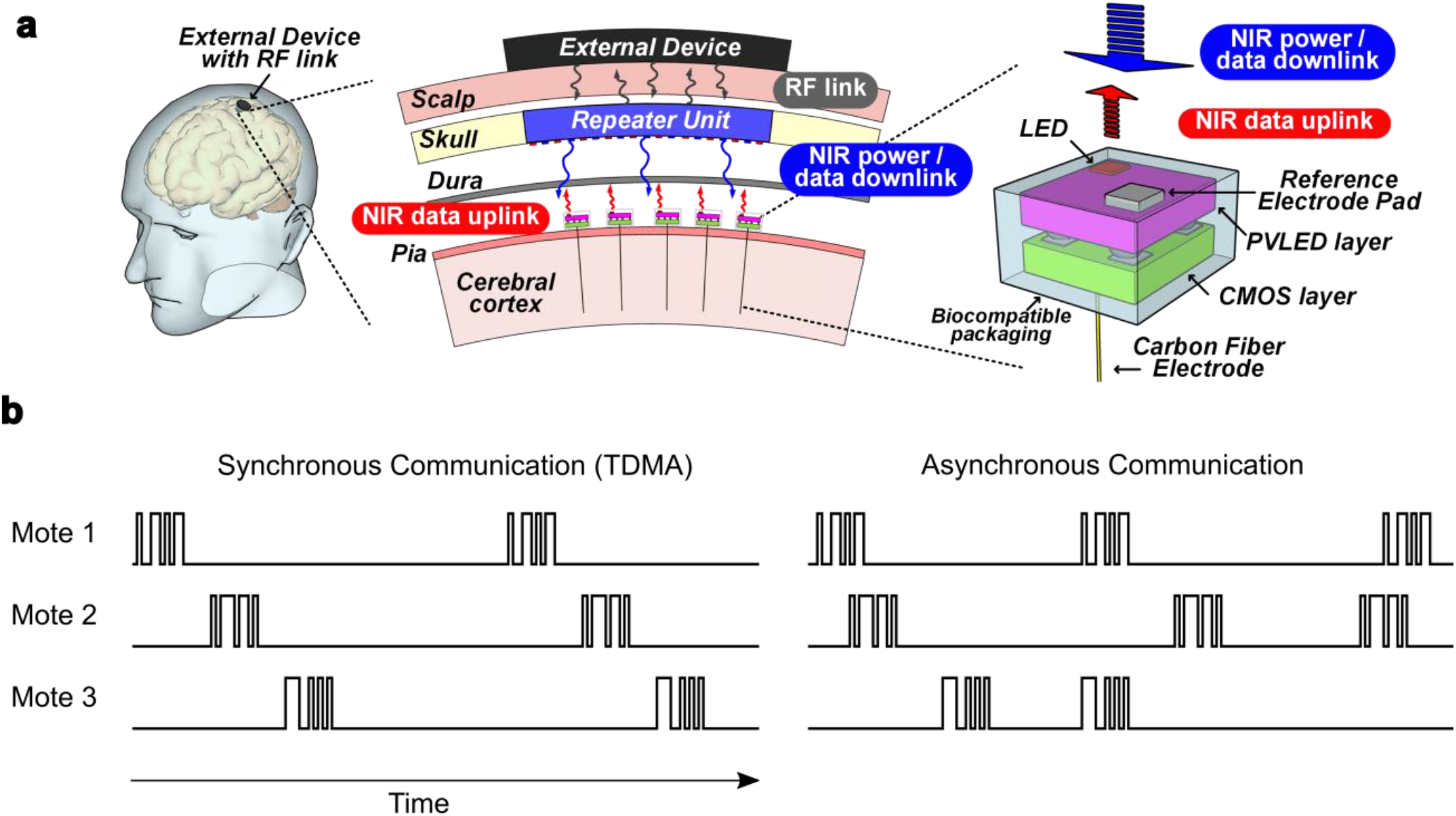
(a) Proposed Michigan Mote system (figure adapted from [23]). Motes sit beneath the dura and are powered by near-infrared (NIR) light. They record cortical signals and transmit data through an IR uplink to a receiver (“repeater”) unit within the skull. The receiver transmits data from all motes to an external device through an RF link. (b) Synchronous vs asynchronous communication schemes. Left – In synchronous schemes like TDMA, each mote has a precise timeslot to transmit data. Right – In asynchronous schemes, motes transmit at irregular intervals, and multiple motes may transmit at the same time.

The total power consumed by each mote comes from sampling, filtering, signal processing, wireless communication, and maintaining a clock; the wireless communication scheme affects all of these parts and is important to optimize. An IR communication scheme uses an optical link with the mote as a transmitter and an array of detectors as the receiver, as shown in Figure 1a. Power use for wireless communications presents a demanding requirement for the mote power budget, where the IR approach can be efficiently scaled using light pulses from a microscale light emitting diode on the mote and high-sensitivity photon counting at the receiver. One relatively simple approach for communicating with hundreds to thousands of devices is time-division multiple access (TDMA) where each device has a specific time-slot to transmit data (Figure 1b). TDMA is used in cell phone communications as well as in the Neurograin system [21]. However, when scaling to 1000s of devices, the on-chip clock rate must be scaled to the megahertz range to support precise transmission timings [21]. While megahertz clocks are feasible for power modalities like RF, they consume too much power for IR (see results). Other communication schemes like frequency division multiple access (FDMA) or code division multiple access (CDMA) are challenging to implement with IR light.

Our group has proposed an alternative communication scheme for IR-based communications, pulse-interval modulation (PIM), in which neural data is encoded by the transmission rate of data packets. PIM is asynchronous (Figure 1b), meaning motes would not have specific transmission times, and could then function with clock speeds on the order of 10 kHz regardless of the number of devices [23]. Since PIM encodes neural data in the analog domain, power-hungry analog-to-digital converters are not required. Finally, individual mote power usage and signal fidelity can be adjusted by changing the average packet rate. Specifically, PIM can encode neural features like Spiking Band Power (SBP), or the signal power in 300-1000 Hz frequency band, which our group has previously shown is dominated by single-units, highly correlated with firing rate, and a high-performance BMI input feature [27]. Using SBP allows the sampling rate to be reduced to 2 kSps (when digitally sampling) for significantly reduced power consumption compared to the 10-30 kHz required for traditional BMIs based on recordings of the threshold-crossing signal. Unlike threshold-crossings, SBP does not require setting a channel-specific threshold, or any other commands to be processed by the device, reducing circuitry and communication needed for thresholds. We previously published the integrated-chip design of PIM-motes [23], [24], but we did not investigate performance using significantly lower data-rates in a closed-loop BMI or the feasibility of receiver complexity.

Here, we aimed to determine if IR-based PIM communication, a scheme without precisely clocked transmissions, could support a high channel count BMI in terms of real-time performance and signal recovery within power limits. By simulating PIM communication in a real-time BMI with non-human primates (NHPs), we found that performance matched the state-of-the-art at data rates of only 100 packets/s (1.3 kbit/s/channel). We found that communication schemes must consider overall lag, since lags as small as 30 ms had a significant performance effect. Additional circuit simulations suggest that, unlike other communication schemes, PIM motes can stay below IR power limits, and despite the complexity of an asynchronous scheme, a receiver could accurately decode data from 1000 motes using 3 mW. Therefore, the receiver power consumption could be of similar magnitude to wired recording systems [28] and is unlikely to be a bottleneck for the mote system.

## Methods

### Microelectrode Array Implants

We implanted two male rhesus macaques (Monkeys N and W, as in [29]) with Utah microelectrode arrays (Blackrock Microsystems) in primary motor cortex (M1) using the arcuate sulcus as an anatomic landmark for hand area, as described previously [30], [31]. In each animal, a subset of the 96-channels in M1, with threshold crossings morphologically consistent with action potentials, were used for offline recordings and closed-loop BMI control. Surgical procedures were performed in compliance with NIH guidelines as well as the University of Michigan’s Institutional Animal Care & Use Committee and Unit for Laboratory Animal Medicine.

### Behavioral Task

We trained Monkeys N and W to acquire virtual targets with virtual fingers by moving their physical fingers (as described in [29], [31]). During all sessions, the monkeys sat in a shielded chamber with their left arms fixed at their sides, flexed at 90° at the elbow, with left-hand fingers placed in a manipulandum. Finger movements were measured by flex sensors (FS-L-0073-103-ST, Spectra Symbol) attached to both doors of the manipulandum and position measurements were recorded by a computer running real-time xPC Target. Flex sensors were calibrated at the start of each experiment session. The computer monitor directly in front of the monkey displayed a virtual monkey hand model (MusculoSkeletal Modeling Software) controlled by the xPC Target computer. In hand control, the virtual hand mirrored the monkey’s hand movements, whereas in brain control the virtual hand was controlled by the decoder.

Each trial began with a spherical target(s) appearing along the path of the virtual finger(s) of interest, where each target occupied 15% of the full arc of motion of the virtual finger(s). For a successful trial, the monkey was required to move its fingers such that the corresponding virtual finger(s) of interest moved into the target(s) and hold its position for 500 ms. On successful trial completion, the monkeys received a juice reward. For closed-loop decoding, the targets were presented in a center-out pattern, as detailed previously [31]. Monkey N performed a 2 degree-of-freedom task (controlling index and middle-ring-small (MRS) finger groups; [29]) while Monkey W performed a 1 degree-of-freedom task (all fingers move together).

### Pulse-Interval Modulation (PIM)

In this work we evaluated simulated-PIM communication in offline and online (real-time) analyses. PIM encodes information by modulating the interval between transmitted data packets. In this application, SBP, or the power in the 300-1000 Hz band, is encoded such that the interval between packets is proportional to 1 / SBP; as SBP increases the interval shortens, and as SBP decreases the interval lengthens (Figure 2). In a hardware implementation, PIM sends the mote’s unique 5-12-bit device identifier within each packet. With PIM, data is encoded by time-intervals rather than transmitted bits, making it extremely bit-efficient and useful for ultra-low-power applications including deep-space satellite communications [32]. By adjusting the average packet-rate one can control the signal reconstruction accuracy for a given time scale; as the packet rate increases, the full-bandwidth signal can be reconstructed with increasing fidelity. Consequently, higher packet-rates allow for better reconstruction of the SBP signal but require more transmission power. The calculations for driving PIM signals can be efficiently implemented on chip through a signal integrator and comparator, sending a data packet once a threshold is reached. Adjusting this threshold adjusts the packet rate [23] (Figure 2). Therefore, on SBP-modulated PIM motes, one threshold parameter would need to be set in contrast to a threshold-crossing-modulated PIM mote which would require two threshold parameters.

**Figure 2:**
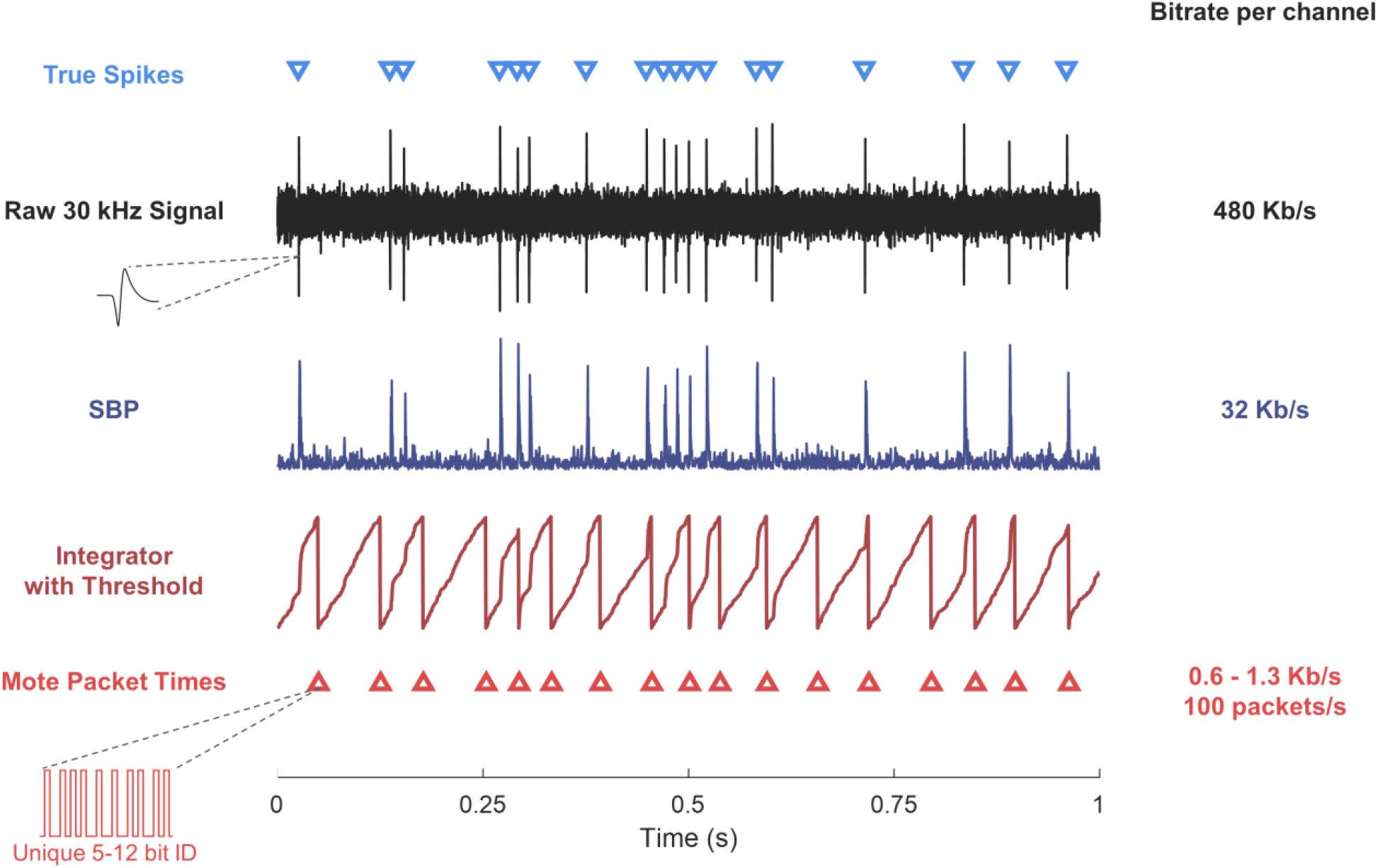
Overview of PIM communication (simulated data). The SBP signal (purple trace) captures much of the same spiking information as the raw 30 kHz signal (black trace) despite a much lower data rate. PIM integrates and thresholds the SBP signal (dark red trace) to determine packet transmission times (red triangles), which contain the mote’s unique ID. When SBP is greater, packets are transmitted more frequently.

Typical BMIs decode average neural activity in 10-100 ms bins, making it unnecessary for motes to transmit every recording sample. Instead, the wireless bit-rate can be significantly reduced by only enough data to recover bin-averages rather than individual samples. Thus, PIM-motes use packets of 10-200 packets/s (pps), substantially lower than the traditional sampling rate of SBP at 2 kSps while maintaining SBP’s BMI-related benefits. On the receiving side, for a single channel, the following equation is used to calculate each SBP-PIM bin:

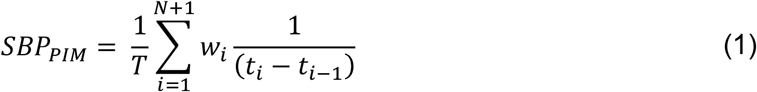

where *T* is the time length of the bin, *N* is the number of packets received within the bin timeframe, *t*_*i*_ is the receiving time of packet *i*, and *w*_*i*_ is the fraction of the packet’s interval contained within the bin (intervals fully within the bin have *w*_*i*_ = 1, whereas the first and last interval represent signal in neighboring bins and have 0 < *w*_*i*_ < 1). The sum is taken over *N* + 1 packets since an additional packet is required to estimate signal within the bin (explained below).

Each PIM packet encodes SBP information from the time preceding the packet arrival; in order to accurately estimate SBP near the end of a bin, the decoder must wait past the end of the bin for a packet to arrive. If a delay is not added, then the end of each bin has an unknown signal value. Thus, when binning PIM packets, a short lag (delay) must be added to account for these packets and improve reconstruction of the true SBP signal (this method was found to better reconstruct SBP bins than without-lag methods). This lag ranges from 15 ms (200 pps) to 50 ms (30 pps), and was determined through offline simulation with real recorded neural data. This extra lag for decoding bins is additive to the BMI’s intrinsic lag (for signal processing and other computational time).

### Signal Processing & Feature Extraction

We recorded 96-channel raw neural data from the monkeys using a Cerebus v1.0 with firmware version 6.03.01.00 (Blackrock Microsystems) digitizing at 2 kSps. First we applied a second-order Butterworth filter to the raw data with a 300-1,000 Hz pass band and extracted the signal magnitude for SBP. Normal SBP bins were calculated by averaging the 2 kSps SBP signal in non-overlapping 16-100 ms bins. Simulated PIM packet times were calculated by integrating and thresholding the 2 kSps SBP signal (with independent thresholds set for each channel). SBP-PIM was then calculated from the packet-times, and binned similarly while accounting for the extra PIM lag (equation 1). For real-time closed-loop decoding, we configured the Cerebus to band-pass filter the raw signals to 300-1,000 Hz. This 2 kSps continuous data was streamed to a computer running xPC Target (MathWorks), which calculated SBP and PIM-SBP in 32 ms bins to be used for decoding.

### Simulated Neuron Recordings

To examine the ability of PIM to accurately transmit neural information at low data-rates, we simulated single neuron recordings and calculated SBP-PIM signals. Using MATLAB (MATLAB 2021a, MathWorks, Natick, MA), we simulated 30 kSps recordings using the method outlined in Nason 2020 (an example trace is shown in Figure 2, black trace). Briefly, a 3 ms averaged sorted unit recorded from Monkey W was randomly placed in time at a desired average spiking rate, and thermal noise was added at the specified SNR. To calculate the SBP signal (Figure 1c, purple trace), we applied a second-order Butterworth filter to the raw 30 kSps signal with a 300–1,000 Hz passband and extracted the signal magnitude. PIM signals were then simulated by integrating the SBP signal and then thresholding the integrated value, where a packet was sent whenever the integrated sum crossed the threshold (Figure 2, red traces). Lastly, to determine how well the SBP and PIM signals captured true firing rate information (Figure 3), PIM pulses were converted to a proxy for SBP, PIM-SBP, by inverting the interval between packets, all signals were smoothed using a 50 ms sliding Gaussian window, and the correlation with true firing rate was calculated.

**Figure 3:**
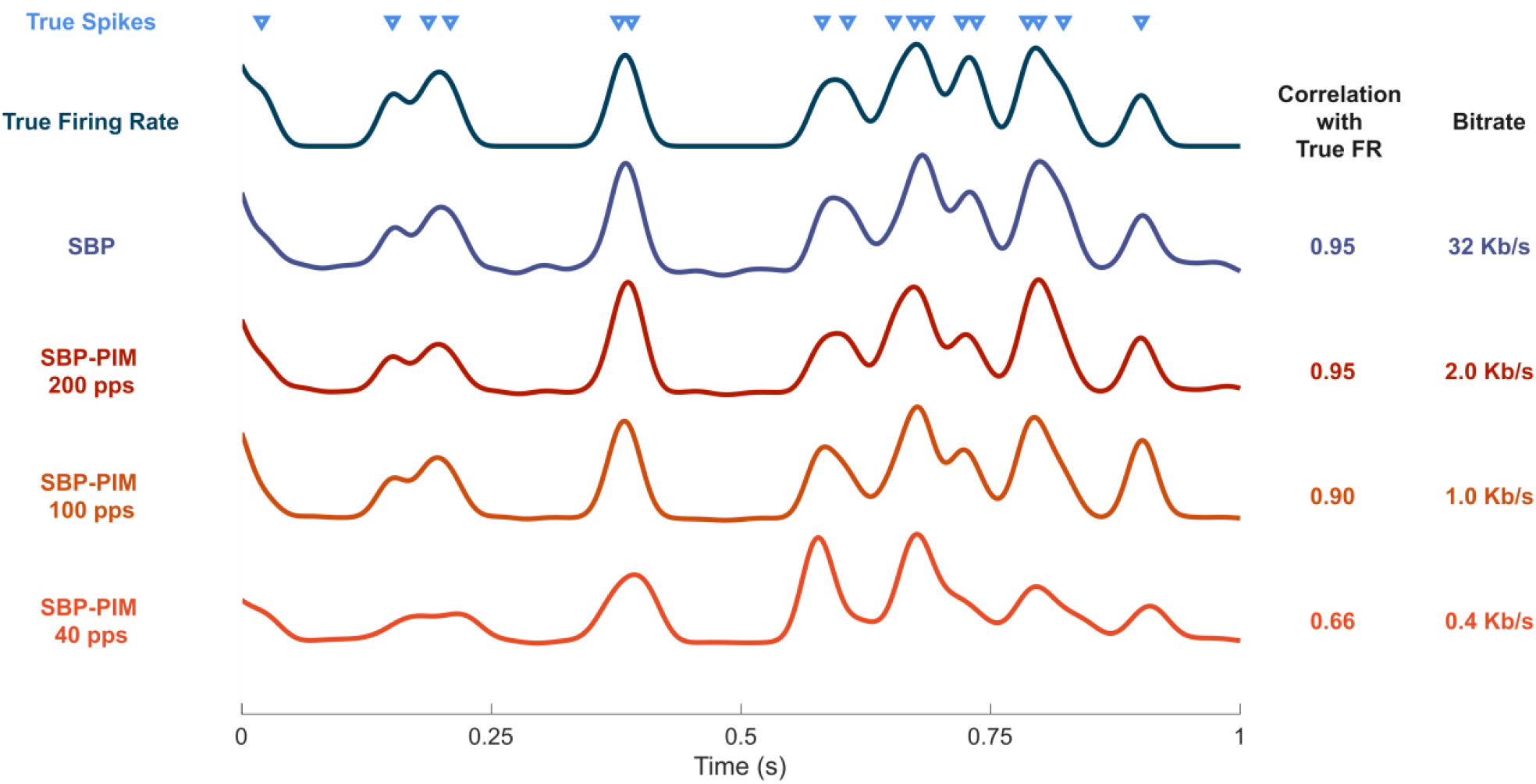
SBP and SBP-PIM correlation with true firing rate. Spikes were simulated and features were smoothed with a 50 ms sliding gaussian window. As the packet rate is dropped, data rate is dropped significantly while correlation with true FR only drops slightly.

### Offline Bin Comparisons

In offline analyses, we compared SBP bins with PIM-SBP bins by calculating the Pearson correlation as well as the variance accounted for (VAF). The VAF was calculated as:

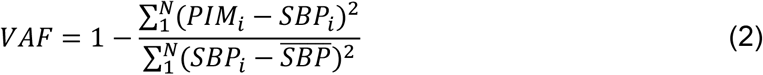

where N is the number of samples (bins), *SBP*_*i*_ is an SBP bin, *PIM*_*i*_ is a PIM bin, and 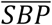 is the mean of SBP bins. Correlations and VAFs were calculated for each channel, and for both metrics a value of 1 indicates that PIM-SBP recovers the same information as SBP.

### Closed-Loop Decoding

For closed-loop decoding, we applied a standard position/velocity Kalman filter with a position/velocity neural tuning model and optimizations as described previously [30] to predict the movements the monkeys made in the behavioral task. On each experiment day, the monkey first performed 400 hand-control trials to use as training data, where hand positions and neural data were simultaneously recorded. Using the training data, separate SBP and SBP-PIM Kalman filters were trained to predict finger positions from neural features in 32 ms bins. Decoders were evaluated during closed-loop brain-control trials.

A total of 13 testing sessions across 6 days were conducted for Monkey N, and a total of 15 sessions across 9 days were conducted for Monkey W. During each testing session, we alternated between SBP and SBP-PIM decoders in an A-B-A-B fashion with approximately 100 trials during individual decoder runs. This alternating method aimed to equalize any performance changes unrelated to the algorithm that occurred within the session. During analysis, the first 10 trials of each run were removed while the monkey adjusted to using the decoder, and the average acquisition-time was taken across all ‘A’ or all ‘B’ trials. Since BMI performance varies across days (due to electrode micromotion, animal motivation, etc.), in some analyses, the SBP-PIM decoder performance was normalized to the average SBP decoder performance to enable cross-day analysis (as in Figure 6).

As previously mentioned, PIM introduces an extra lag not present in the normal SBP decoder, which had a small negative impact on performance. In order to quantify this effect, in some sessions, we added the extra lag to the SBP decoder for an “equalized lag” comparison. While equalizing lag may have slightly lowered SBP performance, this enabled a better comparison of the features. These extra lags are in addition to the inherent lag of 0.5 * bin-size and BMI system lags present in both decoders.

### Feasibility of Synchronous Communication Schemes

To determine if TDMA or other synchronous communication methods would be feasible for a IR mote in terms of power, we simulated the power required for the relatively fast clocks used in these schemes. For TDMA, in a best case scenario, each mote would send an 8-bit sample once every 32 ms. Assuming a 100% overhead for synchronization pulses and guard bits (reasonable due to relatively high clock variability of >1% [21], [23] and similar to the scheme in [21]), the data-rate for 1000 motes is 500 kbps. To account for signal phase synchronization, it is reasonable to require the clock to be 8× faster than the data rate [33], for a best-case (minimum) mote-clock speed of 4 MHz. Then, to estimate power of the on-chip clock, we simulated a ring oscillator in Cadence (Cadence Virtuoso 6.1.7.500.2100 and Cadence Spectre 15.1.0) using TSMC 180 nm CMOS technology. While leaving the mote digital processor connected to the clock generator for realistic output capacitance, we varied clock frequency by adjusting the current flowing through the oscillator, and measured the clock power.

### Relationship between Decode Performance and Receiver Error Rate

To evaluate the effect of receiver error rate on final decode performance, in an offline Kalman filter decode using 96-channel data from Monkey N, we uniformly randomly inserted false-positive packet detections and uniformly randomly removed true packets for false-negative detections. We independently varied the rate of false-positives and false-negatives, and observed the change in finger-velocity correlation and MSE.

### Communication Simulation and Receiver Design

We evaluated the feasibility of PIM communication in a 1000 channel system by simulating the IR optical transmission, detection, and filtering performed by the receiver. In the physical system, motes would be implanted on the cortical surface (beneath the dura) and the receiver unit would be within the skull several millimeters above. Here, we placed 1008 simulated motes in a 28×36 grid with 600 µm pitch, a simulated receiver was placed 2 mm vertically above the motes, and simulated dura mater in-between. All simulations were performed in MATLAB (MATLAB 2021a, MathWorks).

Each simulated mote transmitted at an average rate of 100 pps, where each packet contained 13 pulses (encoding the 12-bit ID through two different intervals corresponding to 0/1 bits).

Packet times were generated by calculating PIM times using real recorded 30 kHz neural data; to generate 1000 channels of data, the 96-channel data was copied with time-shifts to randomize across time. A small jitter of 16 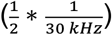 was added to packet times to better simulate the variation in mote clock synchronization.

The simulated receiver used an array of single photon avalanche diodes (SPADs) to detect the mote light pulses. The total number of SPADs ranged from approximately 1000-9000 (see optimization below). An example layout of motes and SPADs is shown in Figure 7a and 7b. Upon detecting one or more photons, the SPAD goes into a high state for the specified dead-time before being reset and able to detect photons again. Thus, SPADs are sampled at a rate equal to 1 / dead-time. We simulated dead times of 100-5000 ns, corresponding to sampling rates of 200 kHz - 10 MHz.

To simulate SPAD detections of light pulses, photons were probabilistically received at each SPAD. Figure 7a shows the probability of a SPAD detecting a mote, based off their relative locations. This probability was calculated using the following equations:

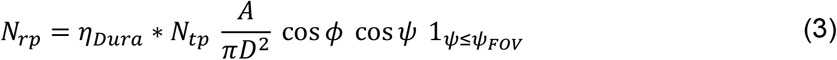

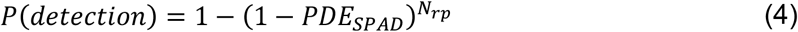

where *N*_*rp*_ is the number of received photons, *N*_*tp*_ is the number of transmitted photons (from the mote source), *A* is the SPAD detector area, *D* is the SPAD-to-source distance, *ϕ* is the radiant angle between source and SPAD, *ψ* is angle between the SPAD normal and source, *ψ*_*FOV*_ is the detector field of view, *η*_*Dura*_ is the estimated transmittance through human dura (0.3, held constant across difference path lengths as a first order approximation) [26], and *PDE*_*SPAD*_ is the photon detection efficiency of the SPAD (approximately 0.1 for 1000 nm light as in [26]).

This optical link model (Eq. 3) is based off [34], with further explanation on its application to motes in [26]. In equation 4, the *P*(*detection*) is the probability of detecting at least one photon in the pulse, which is equal to 1 − *P*(*no detections*), where each detection is considered independent. In addition to this model, SPADs were assumed to have a dark count rate of 1000 counts/s. We assumed the mote plane to be parallel to the SPAD plane, such that *ψ* = *ϕ*. The radius of the circular SPAD detector was set at 30-40 µm such that motes directly underneath a SPAD had >99% probability of detection. Mote light pulses consisted of 1.9×10^6^ photons (assuming an injected charge of 18 pC [23] with an LED quantum efficiency of 1.7% for 1000 nm light [26]).

As seen in Figure 7a, there is relatively wide dispersion of light such that a single mote’s IR pulse may be received by multiple neighboring SPADs with varied probabilities, and each SPAD may receive pulses from multiple motes. As additional motes are added, or the average packet transmission rate is increased, the probability of a SPAD receiving multiple packets simultaneously (a packet “collision”) increases. Thus, the challenge in recovering transmitted mote packets is identifying a given mote’s packet time despite collisions.

To accurately recover packet-times at the simulated receiver, we employed a two-step filter process, as depicted in Figure 7b. First, temporal matched filters were run on each digital SPAD output. Each filter was matched to the ID of a mote; filters performed the convolution of the temporal ID pattern with the input signal. When all signal bits corresponding to known ID times are high, a match is declared. While this detects true packet times with high probability, false-positive matches can occur when light interference is present (for example, the case where the input signal is all high-bits results in a filter match). If a SPAD is local to multiple motes, then separate temporal filters matched to different IDs are run on the same SPAD output through temporal multiplexing.

In the second filter stage, false-positive matches were minimized by combining the filter outputs from multiple local SPADs in a “spatial” filter. The binary temporal filter outputs were weighted by their probability of reception and summed (with 1-bit weights, this is simply the sum of local filter outputs). When this sum was greater than a preset threshold, a match was declared (Figure 7b right and Figure 7c). Finally, accuracy was slightly improved by removing any matches closer than the known ID length (2-4 ms) to another match, since the max true rate of transmission is limited by the ID length.

### Receiver Power Calculation

We estimated the power on the receiver unit required for detecting and filtering transmitted packets in a 1000 mote system, using gate-level power simulations. The estimated total power was taken as the sum of power from running SPADs, reading from and writing to SRAM, running temporal matched filters, and running spatial filters. Each SPAD used an estimated 1.2 nW per detection [35]–[42], where detection rates ranged from approximately 2k to 115k detections/s. As shown in Figure 7b, each SPAD is sampled at 0.4-10 MHz and stored in an SRAM buffer, where SRAM rates are reduced by 16× using 16-bit words for each read or write. SRAM power was calculated using TSMC 28-nm HPC+ technology by summing the power required for read and write operations at a 0.5 V system voltage.

In order to more accurately estimate the power of the non-standard temporal and spatial filter circuits we performed gate-level simulations. The digital filter circuits were designed in Verilog HDL, synthesized using Synopsys DesignCompiler (Version Q-2019.12-SP4) in TSMC 0.028um Logic High Performance Compact Mobile Computing Plus (0.9V/1.8V)(CLN28HT) technology, and simulated using Cadence Xcelium simulator Version 18.03-008, with power estimated using Synopsys PrimeTime PX Version Q-2019.12. Figure 7b summarizes the receiver filter pipeline. Specifically, the input vectors of the gate-level simulations were the SPAD detections of the previously described simulation. Each filter consisted of thirteen 16-bit buffers, where the data in each buffer was read in from SRAM locations corresponding to the preset mote ID times. A 13-bit AND determined if all locations contained high-detections, thus performing the temporal matched filter. 1-5 separate filters were run on the output of each SPAD with different mote IDs, where SRAM read operations were temporally multiplexed. These binary temporal filter outputs were multiplied by 1-bit spatial weights, and summed (initial simulations showed no significant performance difference between 1-bit and higher-bit weights).

### Receiver Power Optimization

To optimize receiver power, we first ran simulations of the following parameter combinations: SPAD pitch from 200-600um, FOV from 45-180°, and sampling rate from 1-10 MHz. For each simulation, we varied the number of temporal filters per mote (the number of local SPADs “listening” for an individual mote) from 0-32, and found the minimum number to maintain less than 1% packet error rate (sum of false-positives and false-negatives). For each number of temporal filters, the spatial filter threshold was optimized to minimize the error rate. Then, given the specific simulation parameters and number of temporal filters per mote, receiver power was calculated. The optimal receiver parameters were chosen as those that achieved the minimum power.

We performed additional simulations with a 30° FOV; with this limited FOV, only 1 mote is visible per SPAD so that interference was eliminated and spatial filter is unnecessary. In these simulations we used a SPAD pitch of 600um, a FOV of 30°, and varied the SPAD diameter from 60-100um, the sampling rate from 0.4-1 MHz, and the number of ID bits from 5-12. The power optimization was performed the same as above.

## Results

### Pulse-interval modulation encodes neural information at low data-rates

We began by investigating how much the data rate could be reduced by using PIM, while maintaining features strongly correlated with the true neural firing rate. First, we simulated single neuron recordings and compared low data rate signals to the true firing rate. Figure 3 shows an example simulated neuron with a firing rate of 20 spikes per second and SNR of 5 (dark blue trace). SBP (light purple trace) was calculated by bandpass filtering, downsampling and taking the signal magnitude of the simulated broadband signal, and the SBP-PIM signals (red/orange traces) were calculated from the SBP signal. Previous work demonstrated that SBP maintains high BMI performance while reducing the wireless data rate from 480 Kbps to 32 Kbps (Nason 2020). In this example, SBP clearly follows the same trend as the true firing rate, exhibiting a correlation of 0.95. The 200 packets/s (pps) SBP-PIM signal has the same 0.95 correlation with true firing rate, showing that it recovers the firing rate information in the SBP signal with an additional order of magnitude reduction in data rate (2.6 vs 32 Kbps).

Next, we explored how well SBP-PIM could transmit real neural data at low-data rates. Using neural data previously recorded from non-human primates performing a virtual finger task, we varied the packet rate and measured the correlation and VAF between binned SBP-PIM and binned SBP in 32 ms bins. A correlation or VAF of 1 would imply that SBP-PIM perfectly recovered the original SBP signal, and could thus achieve the same BMI performance. As the PIM packet rate drops, the signal resolution at finer timescales drops, reducing the correlation with SBP. As in Figure 4, for packet rates greater than 100 pps the curve is relatively flat, showing little benefit to increasing the data rate above this level. The 100 pps signal has only a small reduction in ability to recover SBP (a bin correlation of 0.96) with a major reduction in data rate (1.3 vs 32 kbps), indicating its potential use as a power-efficient BMI feature. However, at lower packet rates the correlation drops more rapidly (correlation of 0.9 at 45 pps).

**Figure 4:**
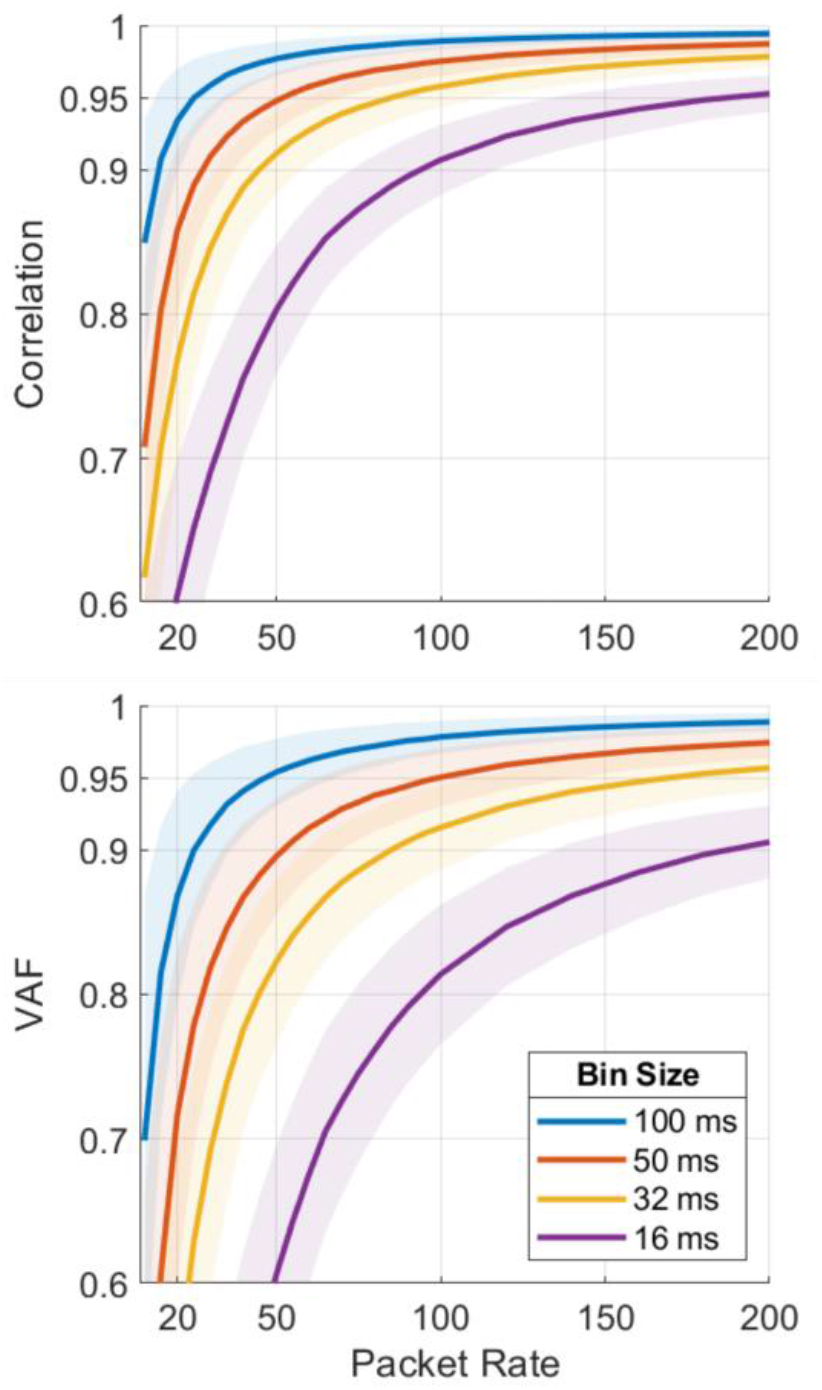
Correlation and VAF between SBP bins and SBP-PIM bins across packet rates and bin sizes. Data is from real neural recordings from Monkeys N and W performing the finger task.

While the previous analyses used 32 ms bins, BMI systems can operate at a range of bin sizes, typically 20-100 ms. Therefore, we also examined SBP vs SBP-PIM correlations for different bin sizes. As seen in Figure 4, longer time bins predictably require less-frequent PIM updates (a lower data rate) to achieve the same correlation with SBP; to achieve a 0.95 correlation, 32 ms bins require 70 pps while 100 ms bins require only 20 pps. In general, for relatively accurate recovery of the original signal (e.g. 0.95 correlation), the packet rate should be roughly twice the bin-rate. This presents a tradeoff between data rate and performance since shorter time bins (on the order of 20 ms) enable significantly higher closed-loop BMI performance [43], [44], but require a higher data rate (and greater power consumption to filter PIM signals).

### Pulse-interval modulation maintains online decoding performance at 100 packets/s

While offline simulations indicate that low PIM data-rates maintain strong correlations to binned features, it is unclear how much signal fidelity is required to maintain performance of a real-time BMI, which incorporates closed-loop user feedback. To evaluate SBP-PIM’s ability to control a real-time closed-loop BMI, Monkeys N and W performed a finger target acquisition task using SBP and SBP-PIM Kalman filter decoders. The SBP-PIM decoder simulated receiving and binning packets in real-time. We started with a PIM packet rate of 100 pps due to its high offline correlation with binned-SBP. During these trials, Monkey N performed a 2-DOF task. Figure 5 shows example trials for each decoder, with solid lines indicating the decoded finger positions and boxes indicating the target. At 100 pps, both decoders had a 99% success rate, where SBP had an average acquisition time of 1.36 sec and SBP-PIM had an average time of 1.26 sec, showing no significant difference *(P > 0*.*05; two-tailed two-sample t-test)*. The 100 pps signal represents a bit rate of 1.3 Kbps. Thus, despite the small loss of information in the offline comparison, the online decoder maintained performance.

**Figure 5:**
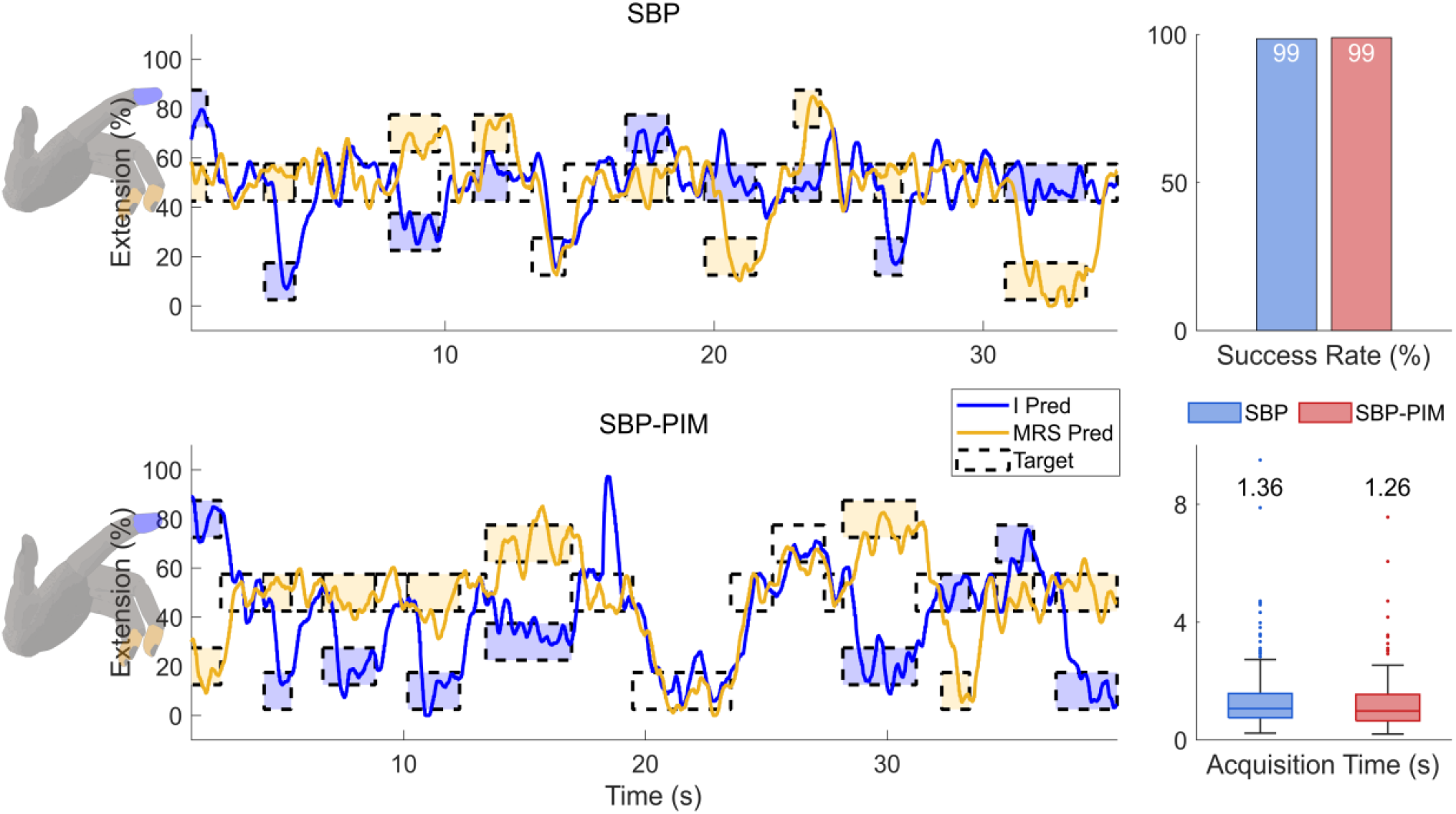
Online 2-DOF decoding with Monkey N using SBP and SBP-PIM decoders. Blue and yellow lines indicate the predicted finger position and boxes indicate the target position. No significant differences for success rate or acquisition time were found (P>0.05; two-tailed two-sample t-test).

To determine how low the PIM data rate could be dropped without closed-loop performance loss, we varied the packet rate from 30-200 pps in separate testing sessions. For the 2-DOF task performed by Monkey N, 100 pps SBP-PIM showed no significant difference in performance compared to SBP on two of three different sessions, as seen by the near zero percent increase in acquisition times in Figure 6a. However, for lower packet rates of 30-80 pps, performance was more than 17% slower with significant differences *(P < 0*.*05 for acquisition times; two-tailed two-sample t-test)*. Monkey W, who had weaker neural signals, only performed a 1-DOF BMI task, and there were no obvious performance trends within this range (tests with 60-150 pps showed with no significant differences on select days, *P > 0*.*05 for acquisition times; two-tailed two-sample t-test)*.

**Figure 6:**
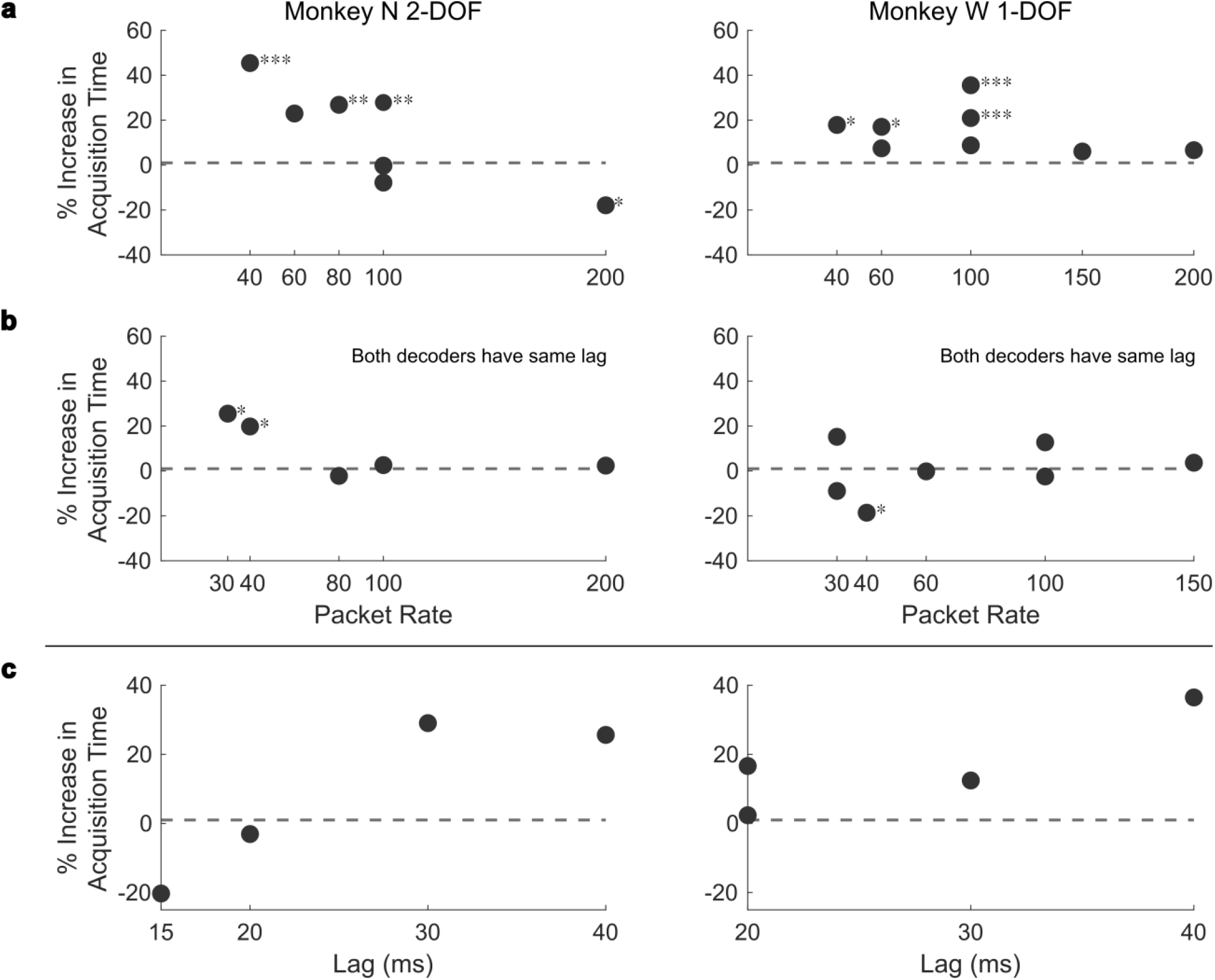
Online SBP-PIM decoder performance. (a) Online PIM-SBP acquisition times as a fraction of SBP acquisition times. In these trials, the PIM-SBP decoder had an additional communication lag not present in the SBP decoder. The dashed 0% line indicates SBP-PIM had the same performance as SBP. Asterisks denote a significant difference from SBP at * P < 0.05, ** P < 0.01, and *** P < 0.001; two-tailed two-sample t-test. (b) Online PIM-SBP acquisition times as a fraction of SBP acquisition times when both decoders had the same lag. The red 0% line indicates SBP-PIM had the same performance as SBP. Asterisks denote a significant difference from SBP at * P < 0.05, ** P < 0.01, and *** P < 0.001; two-tailed two-sample t-test. (c) Increase in acquisition times (AT) from additional lag. The change in AT was calculated by taking the difference in AT between decoders with and without the specified lag.

In these initial comparisons, using SBP-PIM features resulted in an extra delay of 15-50 ms (see methods) before any neural activity would affect the decode due to the asynchronous clocking, whereas the SBP decoder had 0 extra delay. To separate the effects of lag and from feature quality, we added an identical lag to the SBP decoder. In these tests, Monkey N achieved near-equivalent performance for 80 pps or higher *(average acquisition times within 3%; no significant difference: P > 0*.*05; two-tailed two-sample t-test)*, and Monkey W, had similar performance across the full range of 30-150 pps (Figure 6b). Thus, 80 pps PIM might enable equivalent performance if there was not the additional issue of feature lag. This is somewhat surprising, because these results suggest that lags as small as 30 ms have negative performance effects. For example, Monkey N using 80 pps SBP-PIM with a 30 ms lag was 27% slower than SBP, but was 2% faster when lags were equalized (Figures 6a and 6b). Figure 3d summarizes the relationship between increase in acquisition time and lag, with a greater than 12% increase in acquisition times when a 30 ms lag is present. In additional closed-loop tests, we varied the decoder noise level and found that lag may have a greater effect when decoder performance is lower (Supplemental Figure 1).

It is important to note that offline decodes of similar datasets erroneously suggest that lower packet rates could be used without performance loss. An offline velocity Kalman filter using 50 pps SBP-PIM had the same or better VAF compared to an equivalent SBP decoder (Supplemental Figure 2). Thus, offline analyses fail to incorporate the effects of lag, time-averaging due to lower packet rates, and closed-loop control.

### Feasibility of communication schemes at 1000 devices

#### On-device power

An ideal communication scheme would allow for 1000s of motes to communicate with a receiver without bit errors, would stay within system power limits, and would minimize lag. Here, we also considered whether TDMA, a commonly used communication scheme, could be used with IR-powered motes in terms of power consumption. For synchronous communication schemes like TDMA (Figure 1b), the transmitted data-rate and on-chip clock speed must scale to accommodate greater numbers of motes, requiring increased power. To estimate power consumption, we simulated a ring oscillator clock circuit; at 4 MHz, the clock consumed 1.63 µW, which is above the 1.5 µW limit [26] without accounting for other device modules (ADC, filters, etc.) Thus, in this model, TDMA is infeasible for IR-motes. Additionally, TDMA has a lag of 1 time-bin (32 ms) unless a faster clock is used.

PIM, however, is asynchronous and does not necessarily require a faster clock as the number of motes increases. Lim et al. 2021 previously showed that a full PIM-mote can function with less than 1 µW using an 8 kHz clock.

#### Receiver Feasibility

With PIM communication, additional complexity (and power usage) of 1000-device communication is handed off to the receiver unit. To determine if a receiver could stay within power density limits and accurately receive neural signals, we simulated communication with 1000 PIM motes.

Figure 7b shows our proposed receiver design. First, an array of SPADs detect incoming light pulses and each detector output is stored in a local SRAM. SPADs can detect single photons, however, they have a binary output and thus cannot distinguish between multiple motes transmitting simultaneously; with 1000s of motes and wide dispersion of transmitted light (Figure 7a), packet collisions are likely since each SPAD detects an average of over 100k light pulses per second. Next, to resolve these collisions, for one mote, a temporal-filter matched to the mote’s ID is run on the binary outputs of nearby SPADs. If all time samples corresponding to the ID pulse times are high, then the filter output is high. To further reduce the error rate, a spatial-filter combines multiple temporal-filter outputs to reduce packet errors and identify when the mote is transmitting. The receiver simultaneously runs many temporal and spatial filters to recover each mote’s packet times.

In simulation, we used real neural data to generate transmitted mote pulses which were probabilistically received at each SPAD according to the light dispersion model shown in Figure 7a. Using SPADs at 300 µm pitch and a 10 MHz sampling rate, after running temporal matched filters each detector had a 19% error rate on average (1% false-positives, 18% false-negatives; sampling rate of 10 MHz; similar to Figure 7c, black traces). Then, the spatial filters recovered the true transmission times with less than 0.5% error rate (0.1% false-positives, 0.3% false-negatives; 32 filters; similar to Figure 7c, blue trace).

An additional offline simulation involving randomly adding packet errors suggests that offline decode correlation depends on the total error rate (false-positives plus false-negatives). This simulation showed that a 1% error rate corresponded to less than 5% increase in MSE and less than 0.01 drop in correlation for velocity-decoding (Supplemental Figure 3). Thus, for further analyses we aimed to keep the packet error rate below 1%.

Finally, we optimized receiver power consumption by varying SPAD pitch, field-of-view (FOV), and sampling rate. Some tradeoffs emerge when performing this optimization: If the density of SPADs is increased, fewer temporal filters need to be run per SPAD and SPADs are closer to each mote, but the total number of SPADs and SRAM is increased. Another tradeoff is the detector sampling rate (equal to 1 / SPAD dead time); sampling faster allows for finer temporal resolution and fewer false-positive detections after filtering, but filtering and memory operations must be sped up with higher power usage. Additionally, the SRAM size is proportional to the sampling rate since the mote ID is a fixed time length determined by the mote clock and the receiver must store enough samples to capture the full length of a mote ID. With relatively slow mote clocks, IDs are 2.5 ms in length, requiring an 8 kb SRAM per SPAD when sampling at 2 MHz. Figure 7b shows the proposed receiver architecture incorporating SPADs with SRAM memory, followed by temporal and spatial filters.

For each SPAD pitch, FOV, and sampling rate combination, we found the minimum sampling rate and minimum number of temporal filters required to maintain an error rate of less than 1%, and calculated the total receiver power (SPAD power, SRAM power, and filtering power). As seen in Figure 8, FOV had the largest impact on the receiver power. At 180° FOV, all motes are visible to each SPAD, requiring 475 mW to resolve packets (500 µm pitch, 10 MHz sampling rate). However, at 45° FOV, only 5 motes are visible to each SPAD, and receiver power is reduced to 14 mW (600 µm pitch, 2 MHz sampling rate). Across all FOVs, the total power was dominated by the SPADs, accounting for 61-69% of the total.

**Figure 7:**
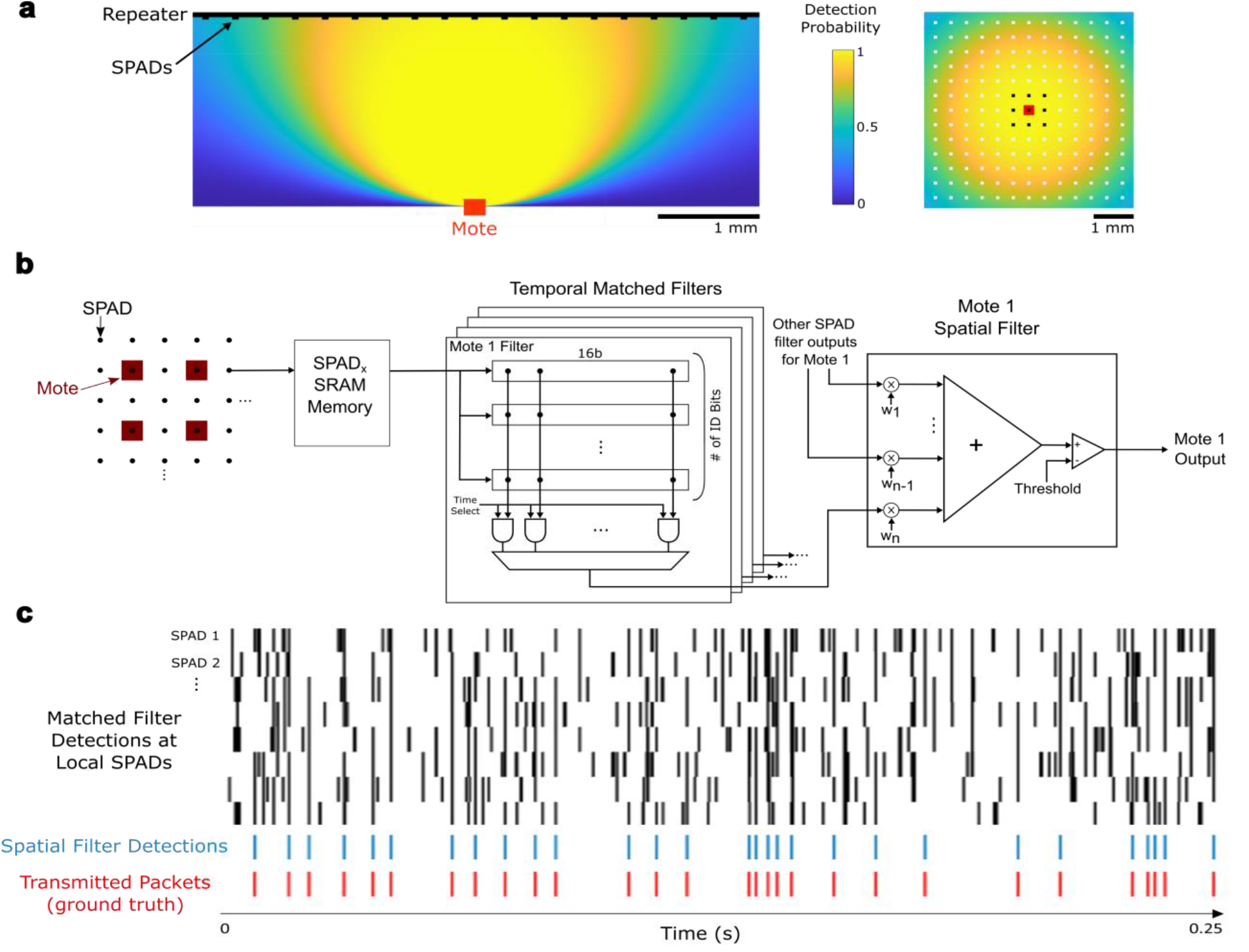
Proposed receiver design and filtering. (a) Probability of IR pulse detection by SPADs. Color indicates the probability of detection. Left – side view with mote in red (transmitting upwards) and receiver in black. Right – top-down view with mote in red and SPADs shown as black and white squares. SPADs in black run temporal-filters to “listen” for the red mote’s ID. (b) Receiver filtering pipeline. Binary SPAD samples are stored in a local SRAM and accessed to perform a temporal matched filter for nearby motes. The outputs from multiple temporal filters are combined in a spatial filter and thresholded to produce the output for each mote. (c) An example simulated detection of one mote using temporal and spatial filters. Black traces indicate the temporal-filter detections at SPADs near the mote. The spatial-filter combines the black traces in a weighted sum and threshold operation to reduce erroneous detections (blue traces), and better match the true packet times (red traces).

**Figure 8:**
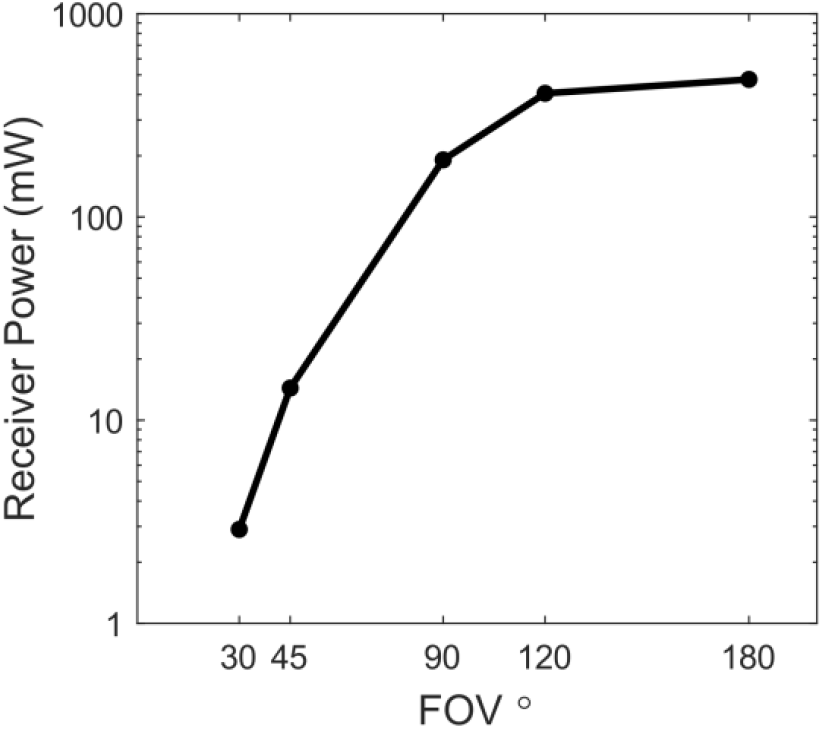
Receiver Power Consumption for less than 1% packet error rate. Each point represents the optimal parameters across all SPAD pitches and at the given FOV.

Once the FOV is reduced to 30° or less, only 1 mote on average is visible to each SPAD, significantly reducing packet collisions and removing the need for the spatial filter. With a 30° FOV, the simulated receiver used 2.9 mW (600 µm SPAD pitch, 80 kHz sampling rate) with an error rate of less than 0.1%, for an additional 5-fold reduction in receiver power. In this configuration, 66% of power was for SPADs, 33% was for filtering, and 1% was for SRAM operations. Furthermore, narrower FOVs would reduce the effect of vertical mote motion on light dispersion, improving receiver robustness. Narrow FOVs could be achieved through using micro-lenses on the motes or SPADs, or by building small walls around each SPAD to reduce light interference.

In addition to the aforementioned processing power, the receiver also provides downlink power to the motes through IR light. To achieve the required 0.73 μW of operating power for each mote [23], given a 190 × 204 µm and 25% efficient photovoltaic cell, and a conservative tissue transmittance of 22%, the receiver needs to produce an illumination intensity of 342 μW/mm2. Given an LED efficiency of 75%, the receiver would use an estimated 456 μW/mm2, or 274 mW for a 2×3 cm craniotomy. Thus, the total receiver power (uplink processing and downlink power) would use approximately 277mW for a density of 462 μW/mm2, largely dominated by the downlink power. This power density is well below the 1.35 mW/mm2 tissue irradiation limit [45], and provides some additional margin for other required circuits.

## Discussion

Here we evaluated a novel pulse-interval modulation (PIM) communication scheme that aims to accommodate thousands of wireless IR neural motes while staying within theoretical power constraints. Through offline simulation, we found that asynchronous intervals can efficiently encode SBP using data rates as low as 100 packets per second (1.3 kbps) and maintain strong correlations to both the true underlying firing rate and the SBP. We then simulated the communication scheme in a real-time 96-channel BMI with non-human primates and found that despite some information loss at 100 pps, online performance matched the state-of-the-art SBP BMI. While PIM enables reduction of mote power usage, the scheme requires increased power and complexity at the receiver unit. Our simulations of the IR communication channel suggest that communication with 1000s of motes is possible with a total receiver power usage of approximately 3 mW. Thus, an IR PIM mote system could stay within safe power density limits for downlink power [26] and for heating due to receiver processing [46].

Surprisingly, we found communication lags as small as 30 ms to have large effects on online, closed-loop BMI performance. The effect of lag is typically lost during offline decoding, unless closed loop control strategies are considered in decoding [47]. Thus, wireless BMIs must consider the lag associated with the communication scheme in addition to other signal processing delays. For TDMA-based communication, lag is dependent on how quickly all motes can be cycled through; if each cycle is 32 ms, then communication lag is 32 ms. For reduced lag, mote clocks must be sped up with stricter precision requirements and increased power usage. In the PIM scheme, however, lag is dependent on the average packet rate, with higher packet rates reducing lag.

These results suggest that current BMI systems, including wired [10], [17], [28] and wireless [21], [48] systems, could achieve similar or better performance with lower data rates and reduced power consumption. While full-fidelity recordings or spike-template matching approaches may be useful for neuroscience contexts, in BMI, performance can be maintained with significantly lower data rates (1 kbps instead of 480 kbps). Mote integrated-chips could achieve state-of-the-art performance using a simple analog front-end with a communication driver, without the need for specific neural-spike detection circuitry.

Additionally, some information loss can be tolerated during online control (similar to the acceptable offline error rate found in [49]). Further information loss could be acceptable with the use of non-linear neural network decoders, allowing for lower data rates and communication power consumption. With greater neural data compression, wireless communication and power transfer is a significantly smaller limitation to the overall BMI. In light-based mote systems, efforts can be put toward optimizing the power of SPAD detectors and developing micro-lens arrays integrated on the receiver [50]–[52].

Wireless mote-based BMIs may enable stable recording for longer time periods than conventional wired arrays. Unlike monolithic recording systems, mote recording-channels could fail independently of each other, allowing for a longer system lifetime. With 1000 or more intracortical recording channels distributed across the cortex, in addition to uncovering novel neuroscience data, there may be enough information to decode many degrees of freedom with high accuracy, setting the path toward clinically effective BMI.

## Acknowledgements

We would like to thank Michael Barrow and Eunseong Moon for assistance in developing the IR light model. We thank Eric Kennedy for animal and experimental support. We thank the University of Michigan Unit for Laboratory Animal Medicine for expert surgical and veterinary care. This work was supported by NSF grant 1926576, the D. Dan and Betty Kahn Foundation grant AWD011321, NSF grant 2129817, and NIH grant 1R21EY029452. J.T.C was supported by NSF GRFP 1841052. S.R.N was supported by NIH Grant F31HD098804. M.S.W. was supported by NIH grant T32NS007222.

## Ethics Statement

This study was carried out in accordance with the recommendations of Guide for the Care and Use of Animals, Office of Laboratory Animal Welfare and the United States Department of Agriculture Animal and Plant Health Inspection Service. The animal care and monitoring protocol was approved by the Institutional Animal Care and Use Committee at the University of Michigan. The study protocol was approved by the Unit for Laboratory Animal Medicine at the University of Michigan.

## Supplemental Information

### Noisy Decoders May Have Increased Sensitivity to Lag

In an additional testing session, we evaluated the effects of lag on decoders with varied noise levels. For these trials, additive gaussian positional noise was added to the decoded finger position. An exponential decay with time constant of 2 seconds was added to the noise to keep the finger position better centered at the true decoded position. The noise standard deviation was set such that the noise was visible and significantly affected performance, but such that targets could still be acquired without decreasing the monkey’s motivation. Variable time lag was also added to the decoder output. After initial hand control training, we collected 250 trials using the standard Kalman filter before recalibrating using the ReFIT technique as described in [29]. The recalibration was performed to achieve maximum baseline performance before lag and noise were added. In the testing session, approximately 100 trials of each configuration were performed with lags in the pattern 0, 10, …, 100, 100, …,10, 0 ms.

As expected, positional noise decreases performance (orange points, Supplemental Figure 1). However, the slope of the acquisition vs lag is slightly steeper for the noisy trials. While this was not tested for significance, this suggests lag may have a greater impact on performance when greater noise is present.

**Supplemental Figure 1:**
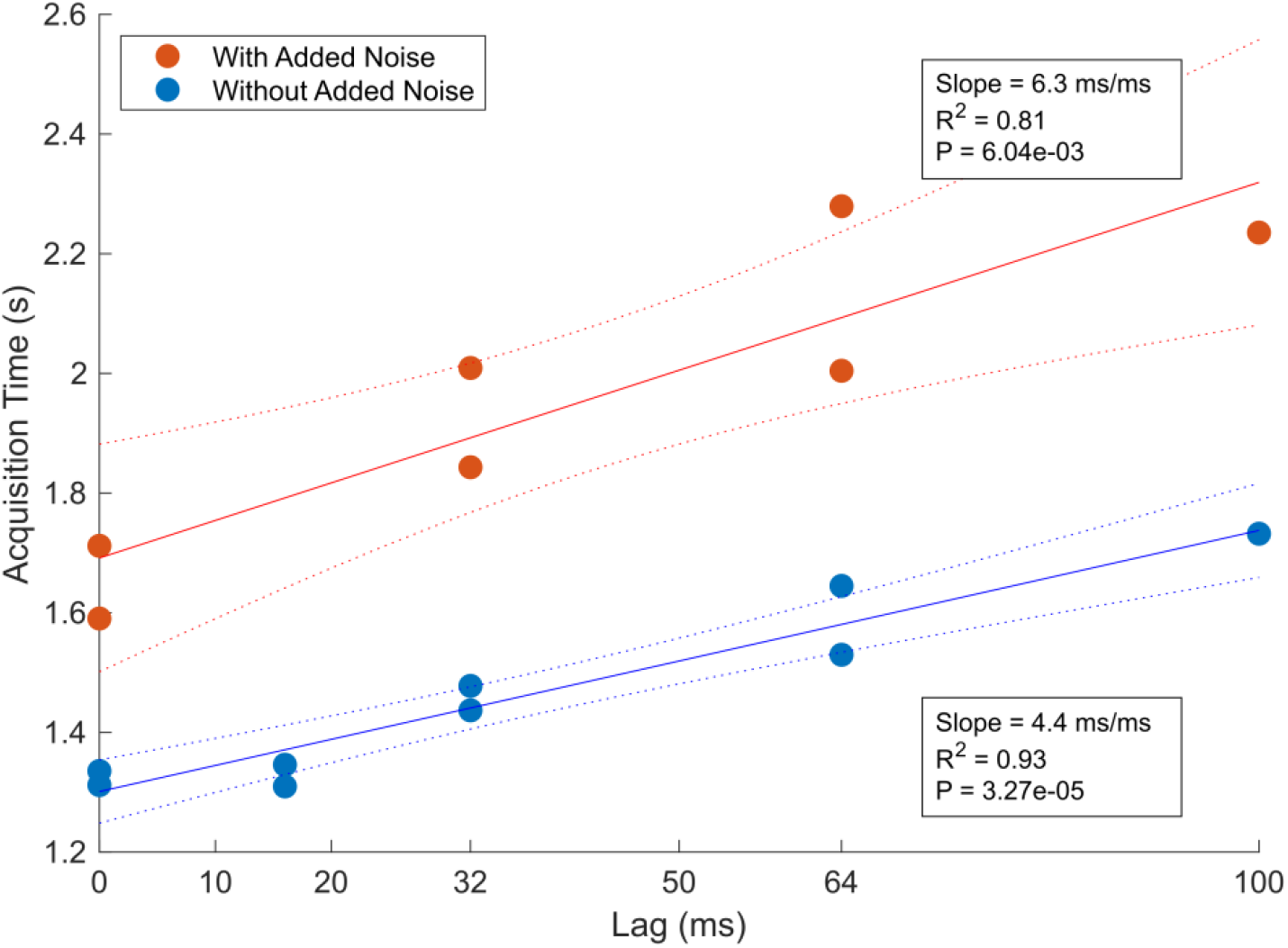
Closed decoder performance for varied lag and positional noise levels. The blue points each represent approximately 100 trials without added noise, with the best fit line in solid blue. The orange points each represent approximately 100 trials where noise was added to the decoded position.

### Offline decodes suggest SBP-PIM maintains performance at low packet-rates

**Supplemental Figure 2:**
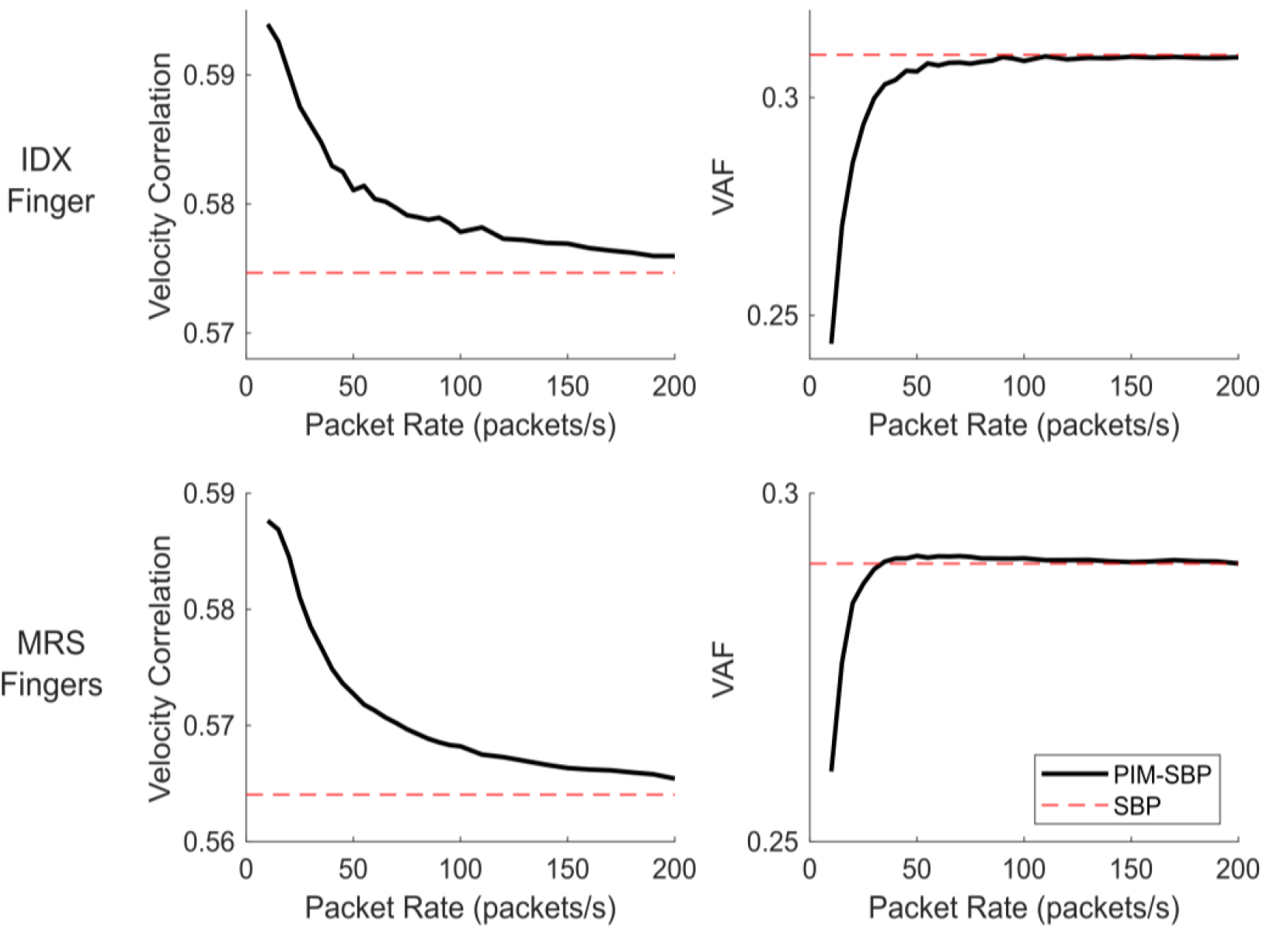
Offline velocity decoding of two-finger groups from Monkey N using a Kalman filter. Black lines indicate the performance of SBP-PIM based decoders compared to the true finger velocities. The dashed red lines indicate the performance of an SBP decoder and represents the baseline performance without simulated-communication effects. Correlations (left) erroneously suggest that packet rates as low as 10 packets/s could achieve higher performance than standard SBP. VAF (right) suggests packet rates lower than 50 packets/s could achieve similar performance to SBP, significantly lower than the results of online, closed-loop trials presented in the main text.

### Offline Decode Performance has a Linear Relationship with Packet Error Rate

To estimate the effects of packet-time errors on velocity decode performance, in an offline analysis, we varied the packet error rate by adding and removing packet-times. In this analysis, we used a single dataset from Monkey N performing the 2-DOF finger task using hand-control. First, we calculated the true packet times for each channel. Next, we uniformly randomly added false packet-times to set the false-positive (FP) rate and uniformly randomly removed true packet-times to set the false-negative (FN) rate. Finally, we used a Kalman filter (Methods) to decode finger velocities, and calculated the correlation and MSE relative to the true finger velocities.

As seen in Supplemental Figure 3a, FP and FN packet errors have approximately the same effect on correlation and increase in MSE. Thus, the sum of FP+FN has an approximately linear relationship with increase in MSE; for a summed error rate of 1%, the percent increase in MSE is limited below 5% (Supplemental Figure 3b).

**Supplemental Figure 3:**
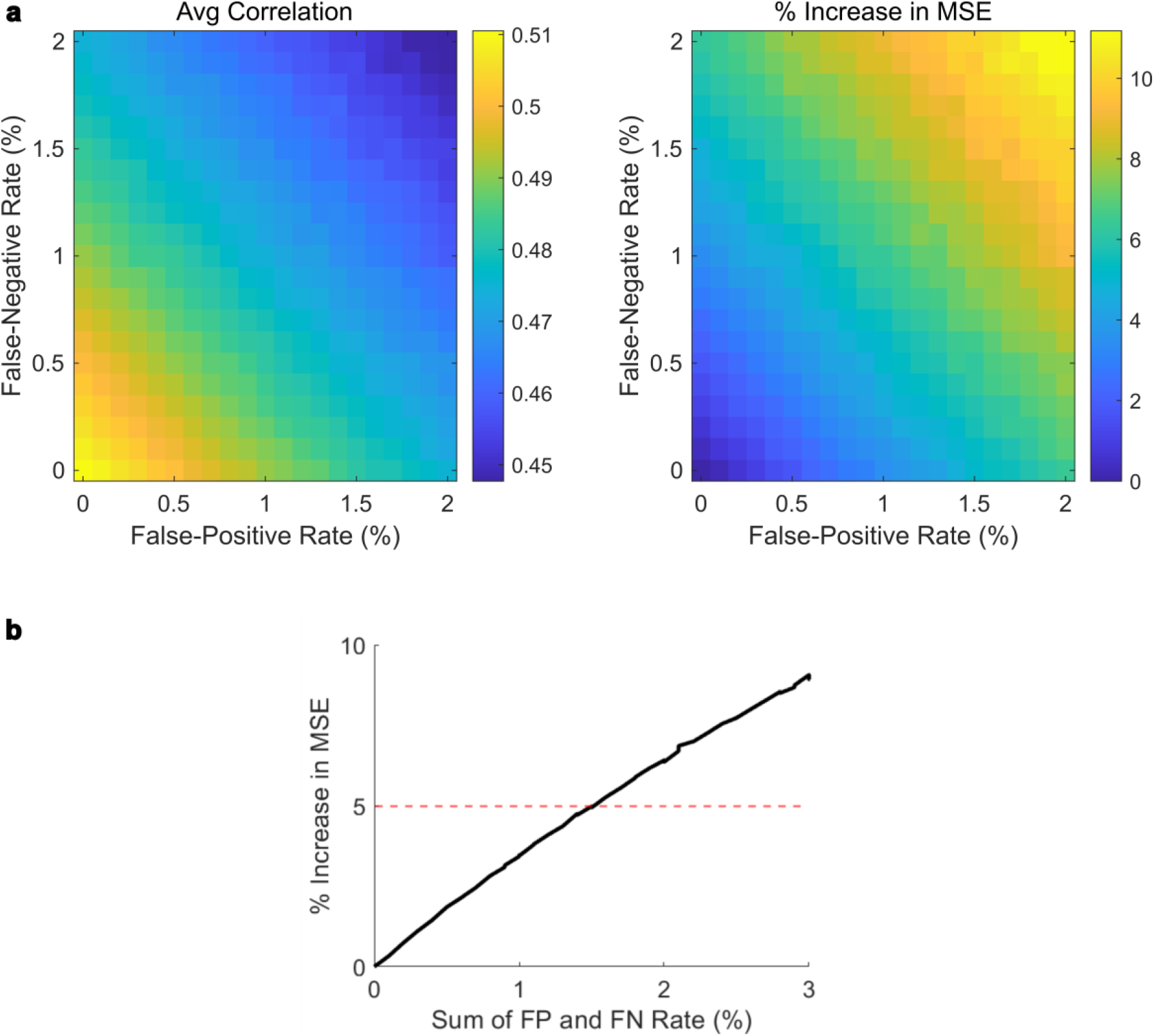
Relationship between offline velocity decode performance and simulated packet error rate. (a) Decode correlations and increase in MSE, where color represents the correlation or MSE compared with true finger velocity. (b) Average increase in MSE as a function of the sum of the FP and FN rate, using the same rates as (a). The red dashed line indicates a 5% increase in MSE, setting an estimated maximum cutoff for packet error rate.

## Notes

### Competing Interest Statement

The authors have declared no competing interest.

